# Geometric differences between nuclear envelopes of Wild-type and Chlamydia trachomatis-infected HeLa cells

**DOI:** 10.1101/2020.04.02.021733

**Authors:** Cefa Karabağ, Martin L. Jones, Christopher J. Peddie, Anne E. Weston, Lucy M. Collinson, Constantino Carlos Reyes-Aldasoro

## Abstract

In this work, the geometrical characteristics of two different types of cells observed with Electron Microscopy were analysed. The nuclear envelope of Wild-type HeLa cells and Chlamydia trachomatis-infected HeLa cells were automatically segmented and then modelled against a spheroid and converted to a two-dimensional surface. Geometric measurements from this surface and the volumetric nuclear envelope were extracted to compare the two types of cells. The measurements included the nuclear volume, the sphericity of the nucleus, its flatness or spikiness. In total 13 different cells were segmented: 7 Wild-type and 6 Chlamydia trachomatis-infected. The cells were statistically different in the following measurements. Wild-type HeLa cells have greater volumes than that of Chlamydia trachomatis-infected HeLa cells and they are more spherical as Jaccard index suggests. Standard deviation (*σ*), and range of values for the nuclear envelope, which shows the distance of the highest peaks and deepest valleys from the spheroid, were also extracted from the modelling against a spheroid and these metrics were used to compare two different data sets in order to draw conclusions.

## 1 Introduction

Electron Microscopy (EM) is an imaging technique that is able to provide a resolving power several orders of magnitude higher than conventional light and fluorescence microscopes and therefore it is ideal to observe very small structures, and of particular interest to this work, of the cellular environment [16, 18]. Modern EM instruments permit the acquisition of contiguous images of the object under observation by slicing very thin sections from the top face of the resin-embedded sample with an ultramicrotome diamond knife [19]. The top face is observed (i.e. an image is acquired), then a slice of the sample is removed and discarded. The sample is then raised up to the imaging position and the imaging starts again. This scanning process continues for a given number of slices, either defined by a user or limited by the thickness of the sample, thus creating a three-dimensional data set. This process is called *Serial blockface scanning EM* (SBF SEM) [8].

The observation of the nuclear envelope (NE) has been a subject of interest for a long time [13, 9]. The NE is a lipid-bilayer membrane, which separates the contents of the nucleus and the chromosomes, from the rest of the cellular structures [5]. The NE contains a large number of membrane proteins with sophisticated roles and functions [21, 6, 12, 10]. The structure and condition of the NE is of great importance as it is related in processes such as cancer [4] or viral infections [11].

Thus, algorithms for the study of the NE are important, in particular the segmentation, visualisation and analysis of the NE could provide parameters to understand the conditions of health and disease of a cell [2, 1, 22, 23, 17, 20].

The processing of EM images is difficult for several reasons. The very large size and resolution of the images, when compared with light and fluorescence microscopy, creates a challenge, as the cells will reveal complex morphological structures. Whilst fluorescence microscopy allows several channels that identify structures of interest, EM only provides a grey scale image and with a reduced contrast between the structures of interest and the background. Thus segmentation is difficult. Furthermore, when serial sections are obtained, the images are transformed into a volumetric data set and it is necessary to link the segmentation of contiguous images to create a final volumetric solution.

Whilst manual delineation of structures, either by an expert or an army of non-experts through crowd-sourcing, is common, this takes considerable amount of time and the results may not always be consistent. Thus, algorithmic solutions that are automatic, consistent and faster than manual processing are preferred.

In this work, we analyse the NE of different types of cells: Wild-type HeLa cells and Chlamydia trachomatis-infected HeLa cells. The volumetric NEs of the cells of the two populations are compared from a geometrical point of view. The NE is automatically segmented and modelled against a spheroid with the algorithms described in [14]. The modelled NE is converted to a 2*D* surface, from which a series of metrics are extracted. The metrics summarise, among other characteristics, the regularity of the NE.

All the code related to this work was performed in programming environment of Matlab^®^ (The Mathworks™, Natick, USA) and is available open source at:

– *https://github.com/reyesaldasoro/Hela-Cell-Segmentation*.

The images are also available from the following repositories:

– **EMPIAR**: *http://dx.doi.org/10.6019/EMPIAR-10094*,
– **Cell Image Library**: *http://cellimagelibrary.org/images/50051*,
– **Cell Image Library**: *http://cellimagelibrary.org/images/50061.*

## 2 Materials and Methods

### 2.1 Wild-type HeLa Cells; Preparation and Acquisition

Details of the cell preparation have been published previously [14], but briefly, the data set consisted of EM images of HeLa cells. HeLa cells were prepared and embedded in Durcupan resin following the method of the National Centre for Microscopy and Imaging Research (NCMIR)[7].

Once the cells were prepared, the samples were imaged using Serial Blockface Scanning Electron Microscopy (SBF SEM) with a 3View2XP (Gatan, Pleasanton, CA) attached to a Sigma VP SEM (Zeiss, Cambridge). The resolution of each image was 8, 192 × 8, 192 pixels corresponding to 10 × 10 nm (Fig. 1a). In total, the sample was sliced 517 times and corresponding images were obtained. The slice separation was 50 nm. The images were acquired with high-bit contrast (16 bit) and after contrast/histogram adjustment, the intensity levels were reduced to 8 bit and therefore the intensity range was [0 – 255]. Then, one cell was manually cropped by selecting its estimated centroid and a volume of 2, 000 × 2, 000 × 300 voxels was selected (Fig. 1d).

**Fig. 1.**
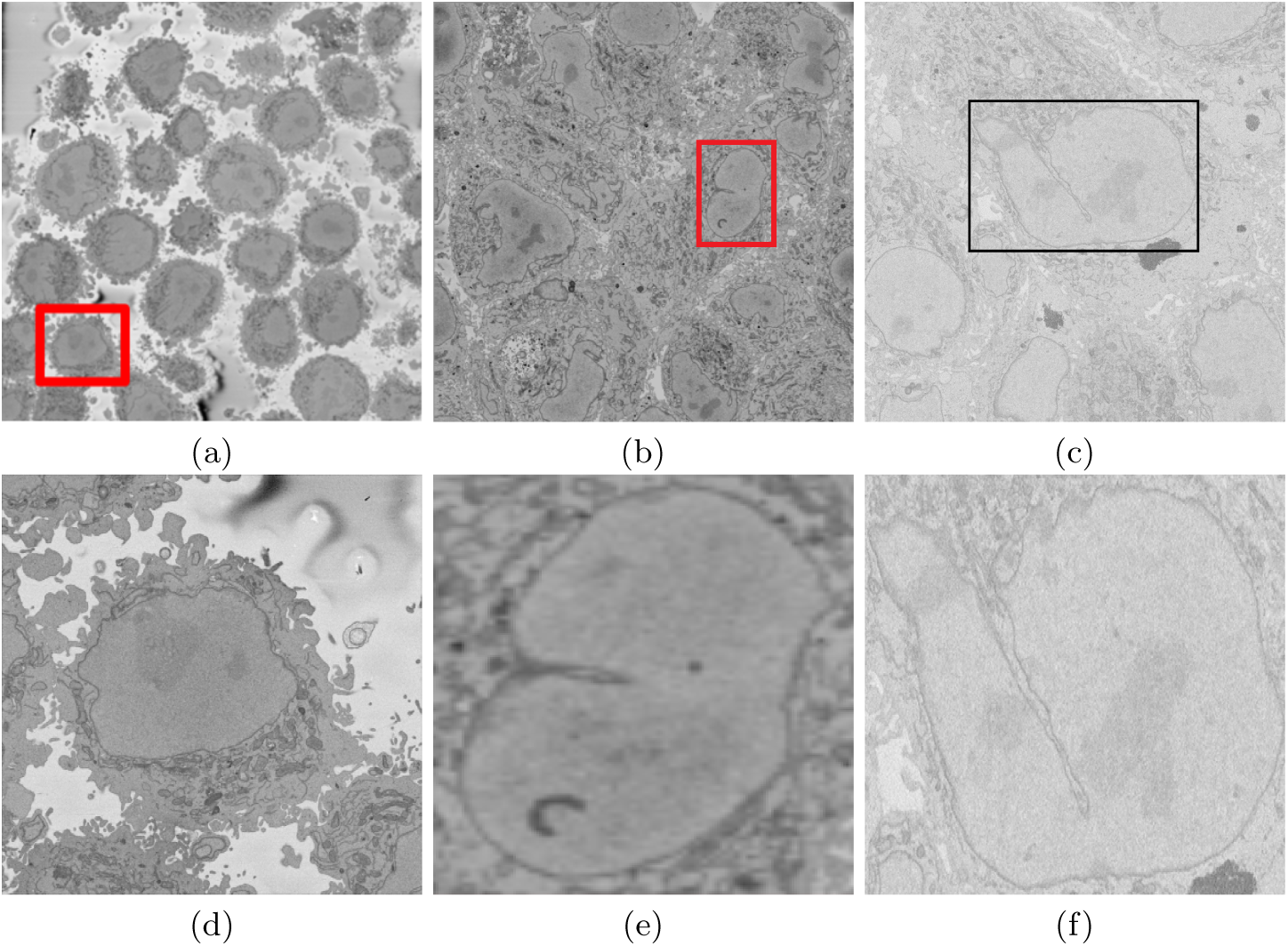
Illustration of the Serial Block Face Scanning Electron Microscope (SBF SEM) images containing Wild-type and monolayers of Chlamydia trachomatis-infected HeLa cells. (a) A representative 8192 × 8192 image from a 3D stack. The Wild-type HeLa cells are the darker regions and the background is a brighter shade of grey. The red box indicates a region of interest (ROI), that is magnified in (d). (b) A representative 3200 × 3200 image from The Cell Image Library (*CIL*50051) arranged as 3D stack (413 slices). Cells have 28 hours post infection (hpi) and voxel size 3.6 × 3.6 × 60 nm. The red box indicates a region of interest (ROI), that is part of a cell which was segmented and modelled against a spheroid in this work. (c) A representative 2435 × 2489 image of the Chlamydia trachomatis-infected HeLa cells from a different data set (CIL50061). Cells in this data set have 12 hpi and voxel size 8.6 × 8.6 × 60 nm. The black box denotes a cell in this slice from this stack (406 slices) that will be posteriorly segmented. (d-f) Detail of the ROIs with a single cell in the centre. The nucleus is the large and fairly uniform region in the centre and it is surrounded by the nuclear envelope (NE) which is darker than the nucleus.

### 2.2 Chlamydia trachomatis-infected HeLa Cells; Preparation and Acquisition

The preparation of the cell has been published previously [15], but briefly, HeLa cells were grown in Advanced DMEM supplemented with 2% fetal bovine serum and 2 mM GlutaMAX-I in 5% CO_2_ at 37°*C*. The cell monolayers were infected with Chlamydia trachomatis serovar L2, strain L2/434/Bu at a multiplicity of infection of 3 in sucrose-phosphate-glutamic acid (SPG). Infections were carried out by centrifugation at 700 × *g* in a Sorvall Legend Mach 1.6 R centrifuge for 1 hour at room temperature. After centrifugation, the inoculum was replaced by fresh cell culture medium and monolayers were incubated at 37°*C* and 5% CO_2_. Chlamydia-infected monolayers were fixed in a solution of 2% paraformaldehyde and 2.5% glutaraldehyde in 0.1 M cacodylate buffer, pH 7.4 for 1 hour. Cells were washed 5X in cold 0.1 M cacodylate buffer then incubated in solution containing 1.5% potassium ferrocyanide and 2% osmium tetroxide supplemented with 2 mM calcium chloride in 0.1 M cacodylate buffer for 30 min on ice. After 5 × 2-min washes in doubled distilled water, cells were incubated in 1% thiocarbohydrazide for 10 min at room temperature. Following 5 × 2-min washes in double distilled water at room temperature, cells were placed in 2% osmium tetroxide in double distilled water for 10 min at room temperature. The cells were rinsed 5 × 2 min with double distilled water at room temperature and subsequently incubated in 2% uranyl acetate at 4°*C* overnight. The next day, cells were washed 5 × 2 min in double distilled water at room temperature and en bloc Walton’s lead aspartate staining was performed for 10 min at 60°*C*. Following 5 × 2-min washes in double distilled water at room temperature, cells were dehydrated using a series of ice-cold graded ethanol solutions and then embedded in Durcupan ACM resin. The resin was allowed to polymerize in a vacuum oven at 60°*C* for 48 hours. SBF SEM imaging was completed using a Gatan automated 3View system (Gatan Inc.) and images were recorded at 60 nm cutting intervals. Two representatives of Chlamydia trachomatis-infected HeLa cell images showing several cells and two in boxes, are shown in Figs. 1b, c and they are magnified in Figs. 1e, f.

### 2.3 Nuclear Envelope Segmentation and Surface Modelling

The methodology for the automated segmentation algorithm has been published before [14], but for completeness is summarised in this section.

#### Segmentation

The images were low-pass filtered with a Gaussian kernel with size *h* = 7 and standard deviation *σ* = 2 to remove high frequency noise. The algorithm then exploited the abrupt discontinuous intensity changes between the NE and the neighbouring cytoplasm (outside the nucleus) and nucleoplasm (inside the nucleus) by applying Canny edge detection [3]. As the NE itself has variations of intensity, there were some cases where the edge corresponding to the NE was broken into several segments. To connect any disjoint segments, these were dilated by calculating a distance map from the edges and then all pixels within a certain distance were included as a single edge. The minimum distance to consider joining was 5 pixels, which could grow according to the standard deviation of the Canny edge detector, which is an input parameter of the algorithm.

The regions of pixels not covered by the dilated edges were labelled to create a series of superpixels. The superpixel size was not restricted so that large superpixels covered the background and nucleoplasm. Morphological operators were used to: remove regions in contact with the borders of the image, remove small regions, fill holes inside larger regions and close the jagged edges.

The algorithm began by segmenting the central slice of the volumetric cell, which was assumed to be the one in which the nuclear region would be centrally positioned and have the largest diameter. Then, the algorithm exploited the volumetric nature of the data by propagating the segmentation of the NE of one slice to the adjacent slice, up and down from the centre. The NE of a previous slice was used to check the connectivity of disjoint regions or separate from the main nuclear region. The algorithm proceeded in both directions and propagated the region labelled as nucleus to decide if a disjoint nuclear region in the neighbouring slices was connected above or below the current slice of analysis. When a segmented nuclear region overlapped with the previous nuclear segmentations, it was maintained, when there was no overlap, it was discarded

#### Surface Modelling

The result of the segmentation of the nuclear envelope is a volumetric surface, with particular shape characteristics, such as notches or invaginations (Fig. 2). To assess the particular geometrical characteristics of these surfaces, a model against a spheroid can be performed. The spheroid was created with the same volume as the nucleus and the position adjusted to fill the NE as closely as possible, as illustrated in Fig. 3a, for one slice and in Fig. 3b for the whole volume where NE is displayed as a red rendered volumetric surface and the spheroid as blue mesh. In order to position the spheroid, the centroid of the segmented cell was calculated, and the coordinates were used as the centre of the spheroid.

**Fig. 2.**
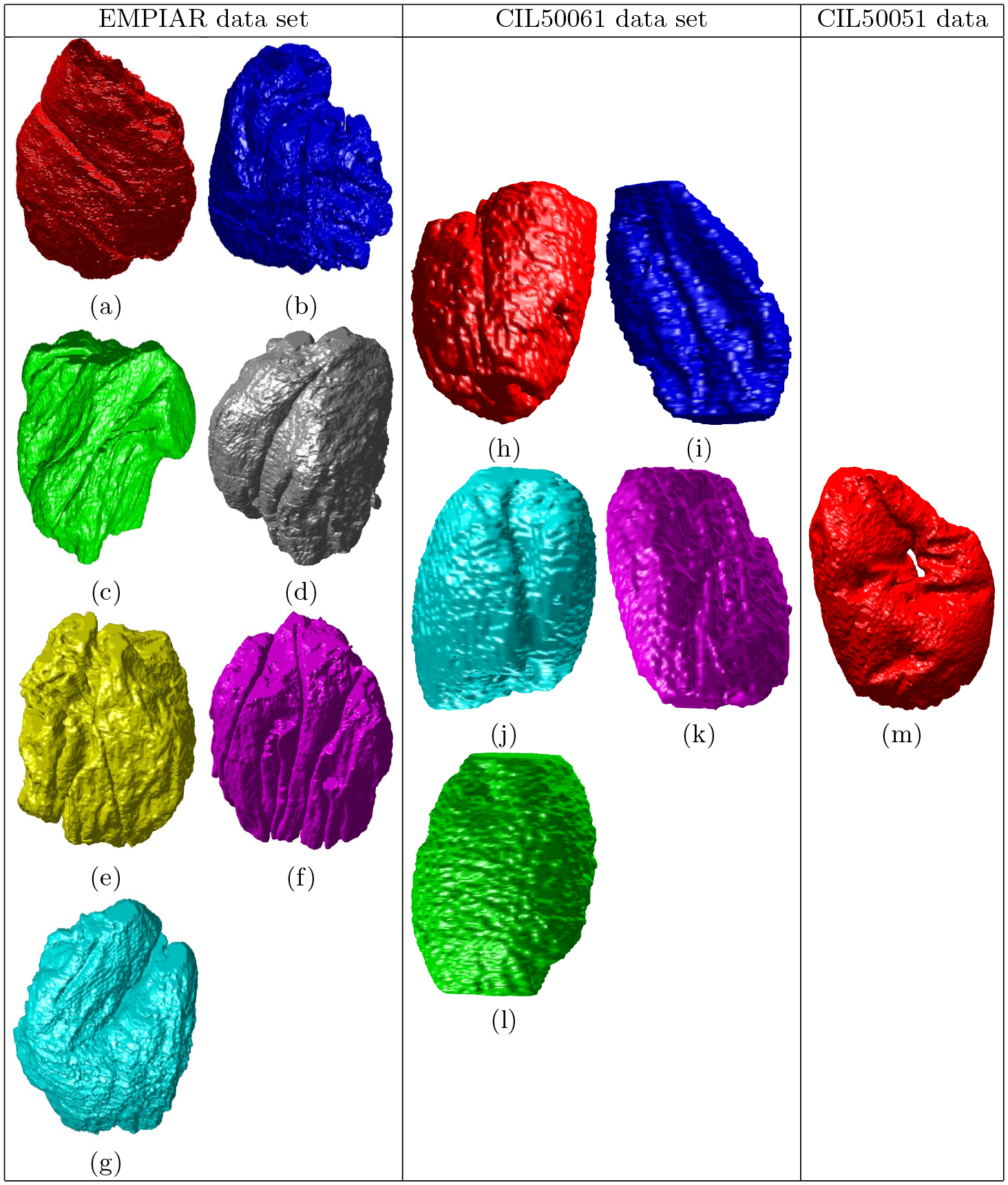
Surface rendering of thirteen Nuclear Envelopes (NEs), different colours are used for visualisation purposes. Seven Wild-type HeLa cells (left column) and six Chlamydia trachomatis-infected HeLa cells from two data sets (centre and right column). (a-g) Wild-type HeLa cells. In each cell, notice the notches that travel up-down along the nuclei (grey (d), yellow (e), purple (f), and cyan (g) cell) and invaginations. Voxel size of Wild-type HeLa cells is 10 × 10 × 50 nm. (h-l) Segmented cells from CIL50061 data set. Cells in this data set are 12 hours post infection (hpi) and voxel size 8.6 × 8.6 × 60 nm. (m) A cell from CIL50051 with 28 hpi and voxel size is 3.6 × 3.6 × 60 nm. Notice the hole of this particular cell.

**Fig. 3.**
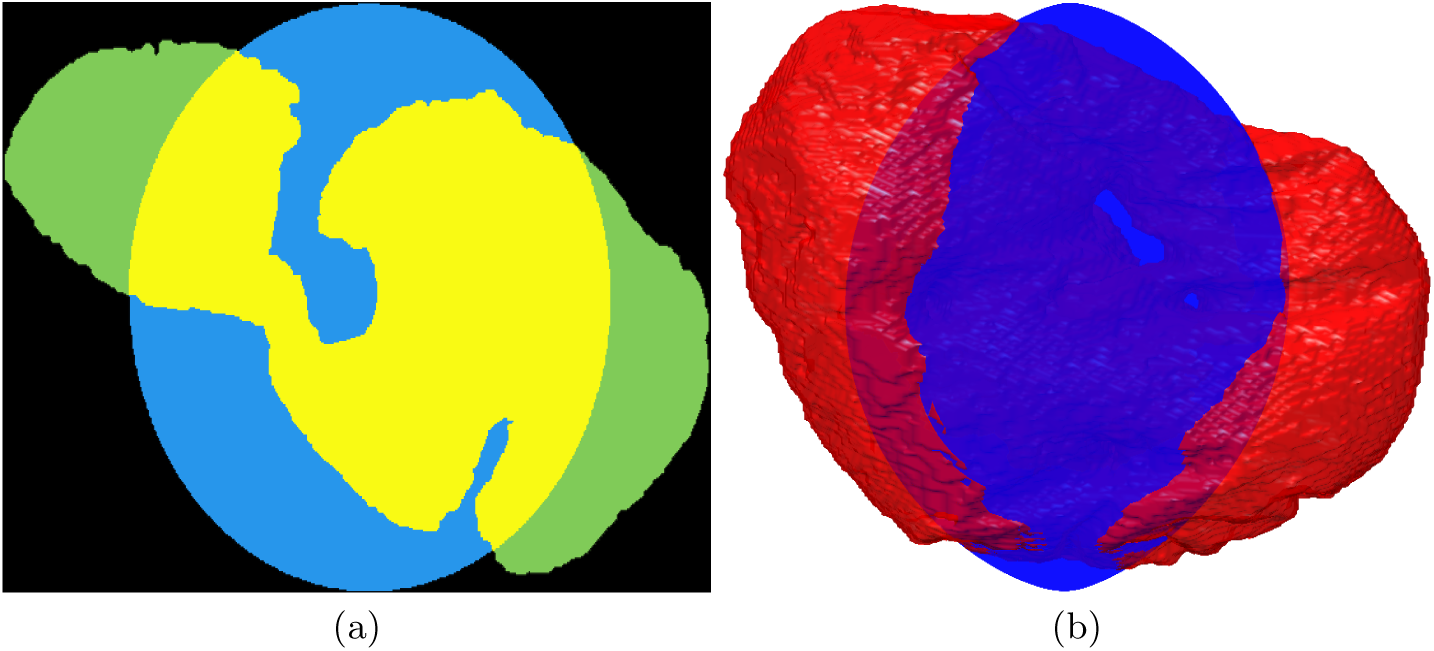
Nuclear envelope (NE) surface modelling against a spheroid. (a) One slice of the modelling where the spheroid is denoted in cyan and the NE in green and the overlap in yellow. (b) Rendering of the NE (red surface) against the model spheroid (blue mesh).

The surfaces of the spheroid and the nucleus were subsequently compared by tracing rays from the centre of the spheroid and the distance between the surfaces for each ray was calculated (Fig. 4a). It was designated that when the NE was further away from the centre, the difference was positive. (Fig. 4b) shows the surface corresponding to the distance from the NE of the first cell to a model spheroid.

**Fig. 4.**
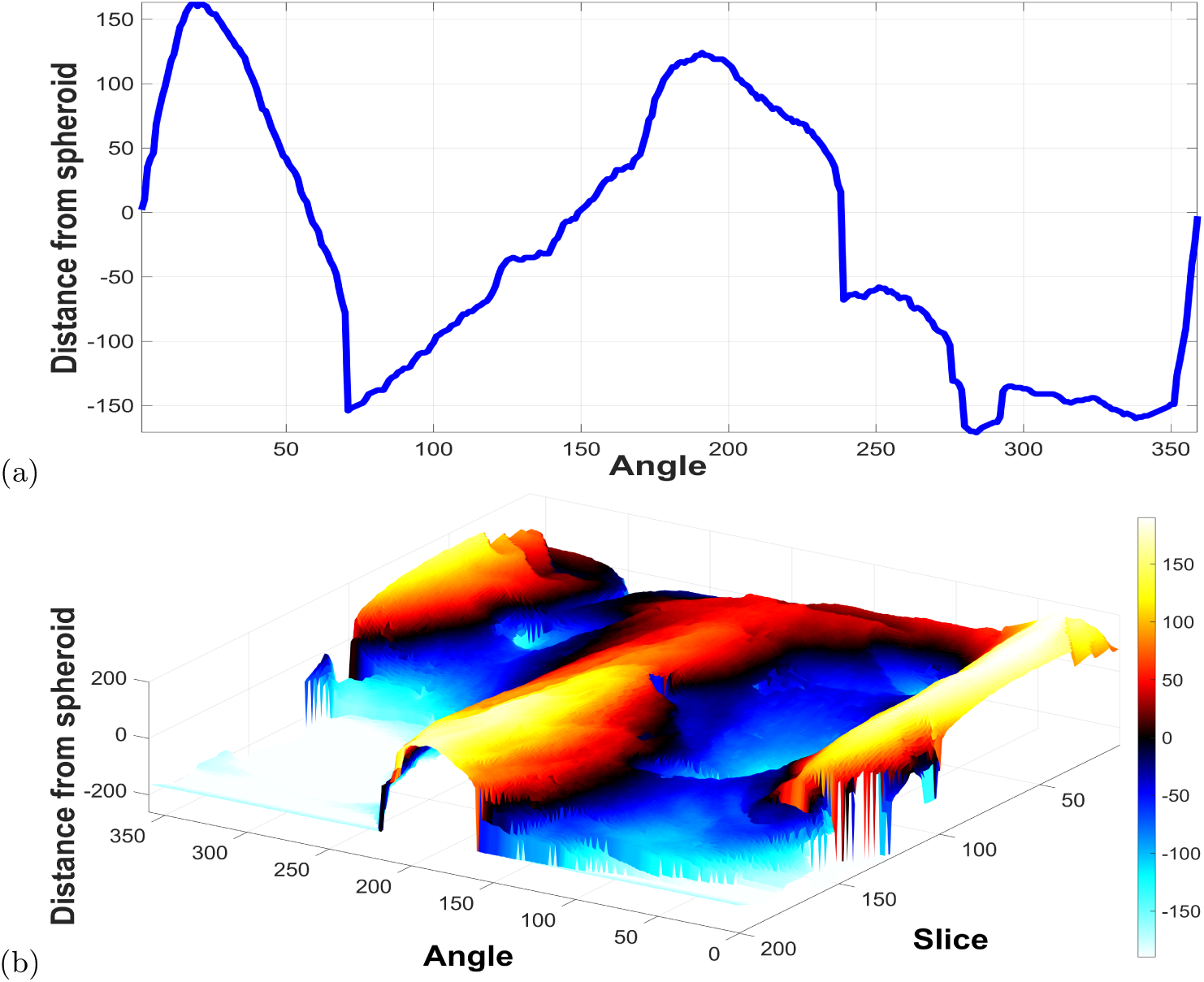
Distances of the NE to the spheroid. (a) Measurements obtained along the boundary of one slice of the NE. (d) Surface corresponding to the distance from the NE to a model spheroid. The surface is formed by placing the lines of each slice as shown in (a).

Several metrics that characterise the NE can be extracted. From the volume itself, the volume of a cell is the first metric to be extracted. To measure the degree similarity between the inner volume of the spheroid and the nucleus, the Jaccard index can be calculated in the following way: the regions where the spheroid and the cell overlap (yellow in Fig. 3a) is considered as True Positive, the region of the spheroid not covered by the cell (blue) is a False Positive and the area of the cell not covered (green) is False Negative. Then, the Jaccard Index is given by:

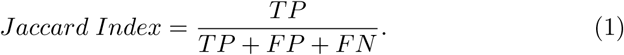

From the surface the following measurements can be extracted from the altitudes of the modelled surface: (1) standard deviation (*σ*) and (2) range of the altitude (distance of the highest peak and deepest valleys from the spheroid).

Finally, the statistical distributions of all volume and surface metrics of Wildtype HeLa cells and Chlamydia trachomatis-infected HeLa cells were combined for comparison.

## 3 Results and Discussion

In this work, an automated image processing algorithm, developed and described earlier in [14], was applied on two different EM data sets, Wild-type HeLa cells and Chlamydia trachomatis-infected HeLa cells, obtained from two different sources. Dimensions of EM images of the data sets were different as well as the voxel size. The algorithm was used to segment and model the volumetric shape of the NE of Wild-type HeLa cells and Chlamydia trachomatis-infected HeLa cells and some conclusions were drawn by comparing results. The NE of thirteen cells, 300 slices each of 7 Wild-type HeLa cells and 406 and 413 slices each of 6 Chlamydia trachomatis-infected HeLa cells, were successfully segmented. Whilst it could be possible to extract more cells from **EMPIAR**, the **Cell Image Library** data sets are restricted as the cells were on the boundaries therefore only six cells were obtained and segmented.

The segmentation of each cropped cell is fully automatic and unsupervised and segments one slice in approximately 8 seconds, one whole cell of Wild-type HeLa cells in approximately 40 minutes. On the other hand, it took almost 60 seconds for the image processing algorithm to segment the whole nuclear envelope of Chlamydia trachomatis-infected HeLa cells. This is for either 406 or 413 slices and an improvement from segmentation and modelling of the Wildtype HeLa cells.

Volumetric measurements revealed that the volume value of the Wild-type HeLa cells is much higher and spread than that of Chlamydia trachomatis-infected HeLa cells (Fig. 5a). This might indicate a shrink in HeLa cells when they are infected. In terms of how close to a spheroid, there does not seem to be a difference between the cells (Fig. 5b).

**Fig. 5.**
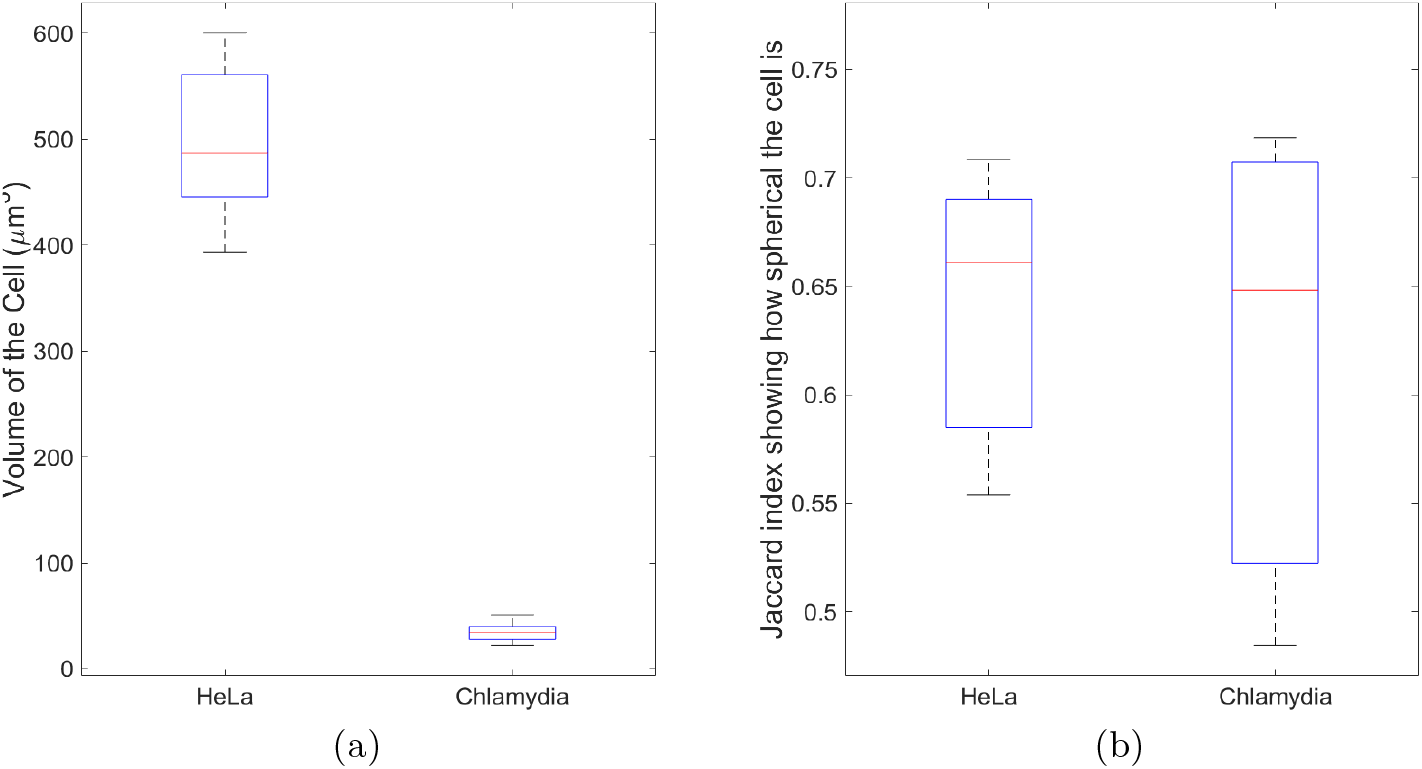
Volume metrics comparison between Wild-type HeLa cells and Chlamydia trachomatis-infected HeLa cells from nuclear envelope (NE) shape modelling. (a) Mean volume value of HeLa cells is much higher than that of Chlamydia trachomatis-infected HeLa cells and this might indicate a shrink in HeLa cells when they are infected. (b) Jaccard index shows how spherical the cells are when modelling against a spheroid.

Surface metrics comparison revealed that the Wild-type HeLa cells had larger values of (*σ*), and range of values for the NE (distance of the highest peak and deepest valleys from the spheroid) than Chlamydia trachomatis-infected HeLa cells, which indicates that the NE surface of the infected cells are smoother than the Wild-types (Figs. 6a,b).

**Fig. 6.**
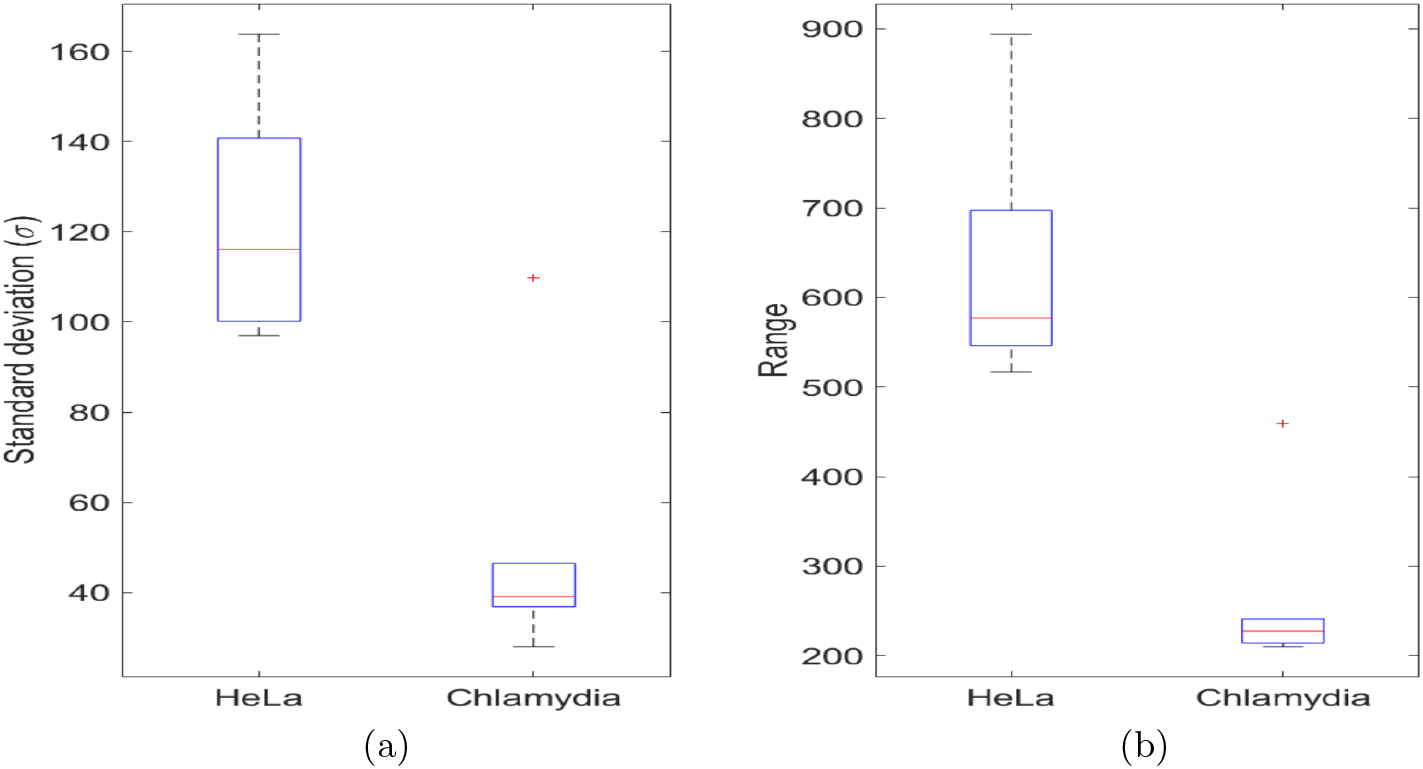
Surface metrics comparison between Wild-type HeLa cells and Chlamydia trachomatis-infected HeLa cells from nuclear envelope (NE) shape modelling. (a) Standard deviation (*σ*), and (b) Range of values for the NE (distance of the highest peak and deepest valleys from the spheroid). These results indicate that the Wild-type cells are far more rugged than the Chlamydia-infected cells.

**Fig. 7.**
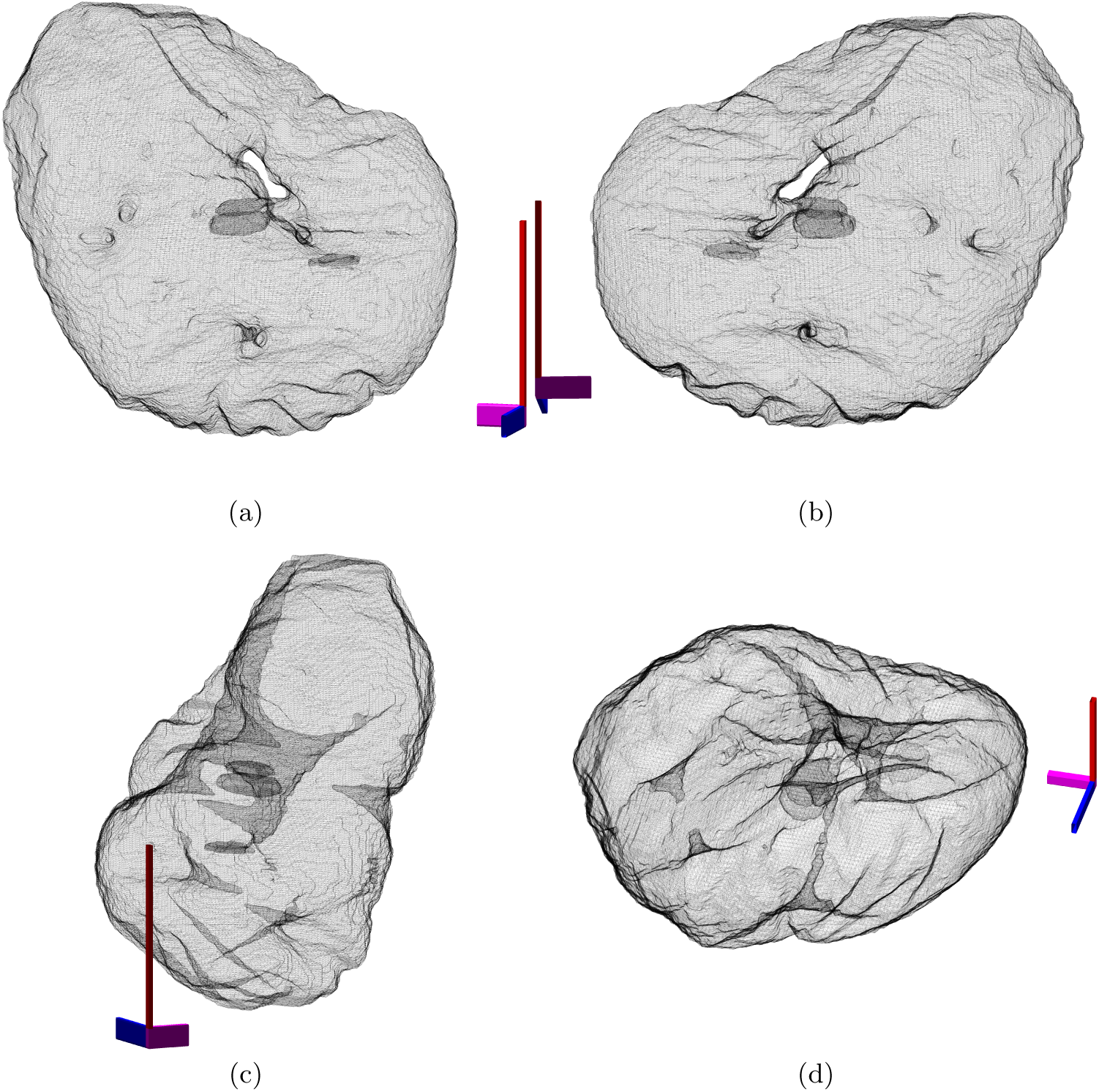
Illustration of the cell from dataset CIL50051. The surface is displayed as a mesh with transparency to show the hole of the nuclear envelope (NE) and the crevices that go deep inside the nucleus. Notice in (c,d) how these invaginations nearly connect separate sides of the NE.

Although the number of cells is relatively low, the results are encouraging and it can be concluded that the Wild-type HeLa cells are geometrically different (smaller, rougher and less smooth) than Chlamydia trachomatis-infected HeLa cells.

One further analysis was performed on the cell from the CIL50051 dataset, the one with hole. Besides the clear hole, the NE has other deep crevices that nearly connect two opposite sides of the NE. This is shown in Fig.7 with the NE rendered with different parameters (no face colour, edges in black and with transparency) and four different view points. Axis are added for reference. Although this level of invaginations and holes were found only in one cell, it is interesting to discover this as it may have significant biological meaning, which is beyond the scope of this publication.

